# GPTAnno: Ontology-tree-guided hierarchical cell type annotation based on GPT models for single-cell data

**DOI:** 10.1101/2025.11.27.690951

**Authors:** Yiran Song, Muyao Tang, Qi Liu, Haofei Wang, Li Qian, Fei Zou, Wenpin Hou

## Abstract

Cell type annotation is critical for interpreting single-cell transcriptomic data but remains challenging due to uncertain cellular clustering granularity and inconsistent labeling across studies. Here we present GPTAnno, an automated, ontology-tree-guided, uncertainty-aware, hierarchical cell type annotation method based on GPT models. GPTAnno directly handles gene expression matrices, integrates multi-resolution clustering with large language model reasoning constrained by the cell ontology to produce standardized, ontology-aware, and reproducible annotations with automatic resolution selection. GPTAnno selects optimal clustering resolutions based on the annotation distance on the ontology tree and quantifies annotation uncertainty to flag ambiguous clusters for expert review. Benchmarking across twelve large-scale datasets demonstrates GPTAnno’s superior accuracy on annotating cell types across various species, tissues, and disease contexts against existing methods. Implemented in R and Python with Seurat and Scanpy compatibility, GPTAnno allows simple inputs to streamline the reproducible annotation, considerably reducing human efforts in repeated reclustering, assigning and examining the labels.

## 1 Introduction

Cell type annotation is a foundational task in single-cell transcriptomics because label quality directly shapes downstream analyses, including comparisons of cell type composition and per-cell-type inferences that inform biology and medicine^1–4^. Determining what constitutes a “cell type” remains nontrivial, as the appropriate granularity is context-dependent^5^.

In many studies, tissues are profiled without prospective sorting and cell identities must be inferred computationally from a gene-by-cell expression matrix using workflows that include normalization, dimensionality reduction, clustering, and marker gene detection, followed by expert curation. Manual curation can be accurate but is time-consuming and requires substantial domain knowledge, motivating automation as datasets and atlases grow.

Existing cell annotation methods fall into three categories. Marker-based methods rely on cellular clustering and expert assignment, referring to established literature^6^ and curated resources such as CellMarker^7,8^, Enrichr^9^, and PanglaoDB^10^. These approaches suffer from ambiguous label granularity across studies, leading to labels at differing levels of detail. When transferring labels to a new study, researchers often assign the majority cell type to a cluster, a process that may require repeated re-clustering and yields limited reproducibility^6^. The conventional marker-based methods require extensive time and human efforts. Reference-based supervised methods (e.g., SingleR^11^, Azimuth^12^, scmap^13^, CellTypist^14^, popV^15^ and AzimuthAPI^16^) can achieve high accuracy when suitable references exist, but they require reference gene expression that are either built into the method or provided by users. In both cases, coverage is typically restricted to a limited set of tissues, constraining the annotatable cell types. More recently, transformer-based approaches (e.g., scBERT^17^, TOSICA^18^, scGPT^19^, scFoundation^20^, Geneformer^21^) have leveraged single-cell data at scale in model pretraining and can be fine-tuned for downstream tasks such as cell type annotation. However, these pretrained foundation models usually require downloading large model weights and often need further retraining or finetuning as new tissue or cell contexts become available, posing barriers to widespread use. In addition, because these approaches rely on supervised classifiers trained on specific label sets that preserve discrepancies across reference datasets, disagreements among methods are common, reflecting differences in label granularity, experiment-specific nuisance factors, and technology-dependent sparsity in the pretrained data^22,23^. Meanwhile, their outputs often lack calibrated uncertainty measures, necessitating substantial human efforts for validation.

Large language models (LLMs) have recently been used to accelerate annotation. Agent-based frameworks, such as CellAgent^24^ and scAgent^25^, enable prompt-based, user-friendly operation and multi-step planning for single-cell analysis, but their annotation component still rely on the aforementioned models (e.g. CellAgent and scAgent leveraged CellMarker) and therefore inherit disadvantages including ambiguous label granularity, inconsistent labels, and a lack of uncertainty estimates. By interpreting marker gene evidence, GPTCelltype^26^ achieved concordance with expert labels across various species and tissues while reducing turnaround to seconds. This strategy has also been incorporated into SpatialAgent’s^27^ CellTypeAnnotation module. However, outputs can be sensitive to clustering resolution and nonstandard label wording unless constrained. In addition, general-purpose LLMs suffers from hallucinations and can generate inconsistent outputs for the same query^28^, underscoring the need for uncertainty measures and robust evaluation. Consensus frameworks and cell-ontology^29^-aware voting have improved robustness and provided useful consistency scores, but they still depend on ambiguous cell type granularity, reference label space and may struggle with emerging types (e.g. popV^15^). While literature mining has been leveraged to enhance performance (e.g., scExtract^30^), reproducibility remains limited by the absence of calibrated uncertainty metrics, necessitating extensive manual scrutiny.

These observations motivate a method that begins from gene expression data rather than only differentially expressed or marker genes, enables data-driven and hierarchical control of label granularity, regularizes label generation against a community ontology, and quantifies uncertainty in reproducible and actionable manner for expert review.

We present GPTAnno, a unified, ontology-tree-guided framework for hierarchical cell type annotation that meets these needs. GPTAnno operates directly on the gene expression matrix, integrates with mainstream analysis ecosystems, and couples data-driven cellular clustering resolution selection with ontology-aware labeling so that parent and child labels remain coherent and standardized across datasets. By combining cross-resolution marker discovery and ontology-constrained reasoning, GPTAnno reduces dependence on differentially expressed gene inputs to GPT models and mitigates sensitivity to *ad hoc* clustering choices. GPTAnno reports uncertainty metrics derived from agreement among candidate labels, margins between top candidates, and ontology-hierarchy consistency, which it uses to suggest the label granularity that yields the most reproducible and ontologically consistence asignments. Users can also leverage these metrics to prioritize ambiguous regions for expert adjudication. To keep knowledge current, GPTAnno harmonizes newly reported markers with the ontology so that definitions can adapt as the literature evolves while maintaining standardized names for cross-study synthesis. With these features, GPTAnno stands out among existing annotation methods (Supplementary Table S1). GPTAnno is released as an open-source package for R and Python, integrates with Seurat^31^ and Scanpy^32^ data structures for practical adoption, and produces uncertainty-aware and ontology-aligned cell type annotation reports.

## 2 Results

### 2.1 Overview of GPTAnno

The workflow of GPTAnno is summarized in Fig.1, which integrates clustering, LLM-based reasoning, and ontology-aware automatic cell type annotation in single-cell and single-nuclei RNA-seq data. Starting from a gene expression matrix, GPTAnno generates biologically context-aware predictions by incorporating tissue information and marker genes, and standardizes annotations by aligning with the Cell Ontology (CL^29^), without requiring any external reference dataset. CL is a controlled vocabulary that standardizes the definition of cell types, including anatomical structures, cell types, and functional roles across species. By encoding hierarchical relationships and linking canonical labels to their synonyms, CL provides a consistent framework for annotation. Mapping the cell type names output from LLMs (OpenAI GPT models) to standard CL terms is a crucial step in GPTAnno, where we can use the CL distance on the ontology tree structure to calculate reliability as one of the criteria to select the optimal clustering resolution. CL has been adopted in major single-cell programs, including The Encyclopedia of DNA Elements 4 (ENCODE^33^), Human BioMolecular Atlas Program^34^, Human Cell Atlas^35^, Expression Atlas^36^, and Kidney Precision Medicine Project^37^, underscores its value for integrative analysis. Standardization through CL is especially critical in large-scale single-cell studies, where heterogeneous naming conventions and the continual discovery of novel subtypes pose challenges for reproducibility and cross-study integration^38^.

**Figure 1.**
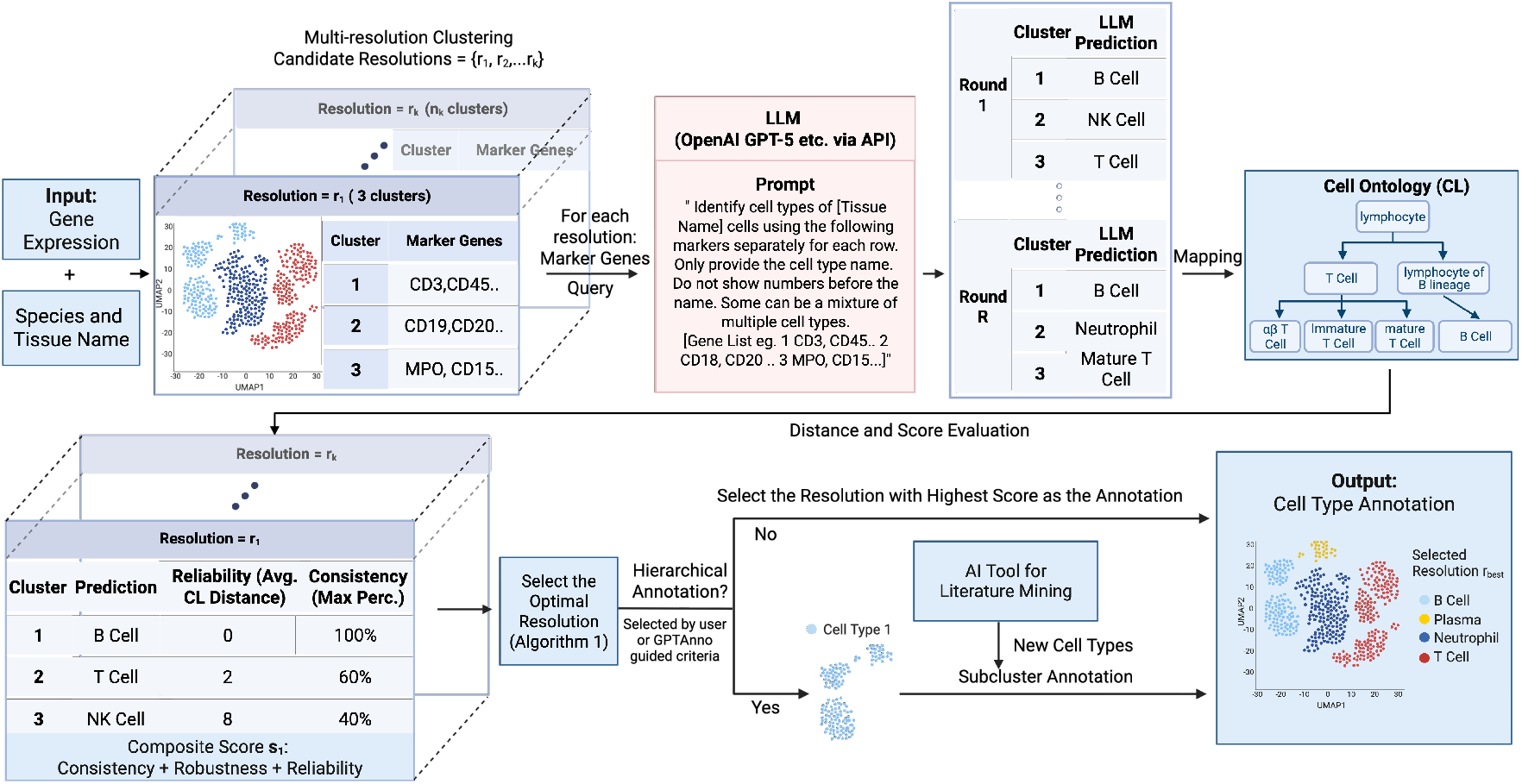
GPTAnno automated workflow for ontology-tree-guided, automatic resolution selection, and hierarchical cell type annotation with uncertainty metrics.

A key strength of GPTAnno is its ability to provide reproducibility measures on CL-grounded GPT annotation and select the most reliable cellular clustering resolution. Rather than relying on a single, subjective cellular clustering choice, GPTAnno evaluates multiple candidate resolutions. For each resolution, differentially expressed marker genes and tissue context are used to prompt GPT across repeated queries. The resulting annotations are then mapped to CL terms to ensure consistent terminology across queries and enable ontology-based evaluation of reproducibility. Three measures are then computed: the average ontology distance (reliability), the average of the maximum reproducible percentages (consistency), and the minimum of the maximum reproducible percentages (robustness, a safeguard against poorly defined clusters). By converting model variability into interpretable distance metrics on the ontology tree, GPTAnno establishes a transparent measure of annotation quality not available in previous methods.

Furthermore, GPTAnno provides hierarchical clustering and subcluster annotation. Specific parent cell types are further subclustered, using modified prompting strategies that either restrict predictions to ontology-defined children or combine subcluster markers with parent markers without restrictions. The first strategy allows GPTAnno to identify canonical subtypes when defined in CL, while the second strategy can discover new cell types reported in new literature but absent from existing CL terms. To further support this process, GPTAnno integrates AI-driven literature mining, incorporating recently described marker genes and cell type definitions into the annotation workflow. Together, GPTAnno transforms cell type annotation from a subjective, *ad hoc* task into a systematic and ontology-grounded workflow that offers reproducible and biologically appropriate annotations.

### 2.2 Evaluation and Comparison with Existing Methods

We systematically benchmarked GPTAnno across 12 datasets from ten studies^39–48^, covering three species (human, mouse, and zebrafish), hundreds of tissue and cell types, developmental and aging stages, and both normal and disease samples such as the samples of cancer and heart injuries. Evaluations were based on GPT-5’s (API version without web access). Although it has been demonstrated that GPT models generalize well to new studies published beyond the models’ training cutoff dates^26^, we carefully include datasets that were published beyond GPT-5’s (API version without web access) training cutoff date Sep 30, 2024.

GPTAnno’s advantages stem from its design across multiple dimensions (Supplementary Table 1), and we demonstrate its superior performance through systematic comparisons with the GPT-based method GPTCelltype^26^, reference-based methods singleR^11^ and Azimuth/AzimuthAPI^12,16^, and the consensus multi-algorithm framework popV^15^. To quantify annotation quality, we introduced a weighted agreement score by classifying prediction–manual pairs into five categories based on the CL hierarchy: exact match (same CL term), child match (prediction is a descendant of the manual label), parent match (prediction is an ancestor), sibling match (prediction and manual label share a common parent within three edges), and no match (unrelated within three edges or not mapped to CL). Weights are assigned as follows: 1.0 for exact and child matches to reward accurate or more detailed predictions consistent with broader labels, 0.5 for parent and sibling to acknowledge partial agreement, and 0 for unmatched terms. This way, we reward biologically coherent predictions while distinguishing true mismatches.

**Table 1.** Domain-specific abbreviation–to–full-name mappings used during cell-type normalization.

| Abbreviation | Full form |
| --- | --- |
| cm | cardiomyocyte |
| cmcs | cardiomyocytes |
| fb | fibroblast |
| ec | endothelial cell |
| epc | epicardial cell |
| smc | smooth muscle cell |
| vsmc | vascular smooth muscle cell |
| endo | endocardial |
| epi | epicardial |
| vasc | vascular |
| atr | atrial |
| vent | ventricular |
| avc | atrioventricular canal |
| oft | outflow tract |

Across the benchmarks, GPTAnno achieved the highest agreement score in all 12 datasets (Fig. 2A) and stood out on non-mammalian data. The lone exception is GTEx, where GPTAnno and popV tie. This is possibly due to the coarse ground-truth granularity and strong reference coverage, which yield clear marker signals for common cell types, pushing both methods to the same parent-level calls. Breaking down match types, GPTAnno shows the highest fractions of Exact and Child matches (i.e., exact or more specific CL terms) and correspondingly fewer Parent/Sibling/No-match outcomes, especially in BCL, Colon, Lung, PDAC_sn (pancreatic ductal adenocarcinoma (PDAC) single-nuclei RNA-seq), PDAC_sc (PDAC single-cell RNA-seq), and zebrafish datasets (chamber and regeneration) (Fig. 2A). PopV (prediction and majority-vote) is often second-best but skews broader (higher Parent matches), reflecting its reference consensus, and it does not apply to non-mammalian species such as zebrafish. GPTCelltype works across all benchmarked species and is competitive on several datasets, but it can trail popV in some settings, underscoring the value of an ontology-based, uncertainty-aware framework like GPTAnno. By contrast, SingleR favors broader Parent labels (e.g., in GTEx and MCA), yielding moderate agreement with reduced specificity. Azimuth/AzimuthAPI are preset-sensitive as broad, medium, or fine. Finer settings raise Sibling/No-match rates when naming diverges from CL, producing uneven accuracy across tissues. This is likely due to its use of DISCO naming conventions^49^, which differ from CL and limit ontology-aware evaluation. A practical advantage of GPTAnno is its species flexibility. Unlike reference-based methods that require pretraining for particular organisms, GPTAnno robustly annotates zebrafish datasets. We also see sensitivity to tissue context in prompts; for example, when annotating mouse aging non-cardiomyocyte hearts^46^, omitting “non-cardiomyocyte” and prompting only “mouse adult heart” led GPTAnno to mislabel a fibroblast subcluster as cardiomyocytes, highlighting the importance of explicit tissue context.

**Figure 2.**
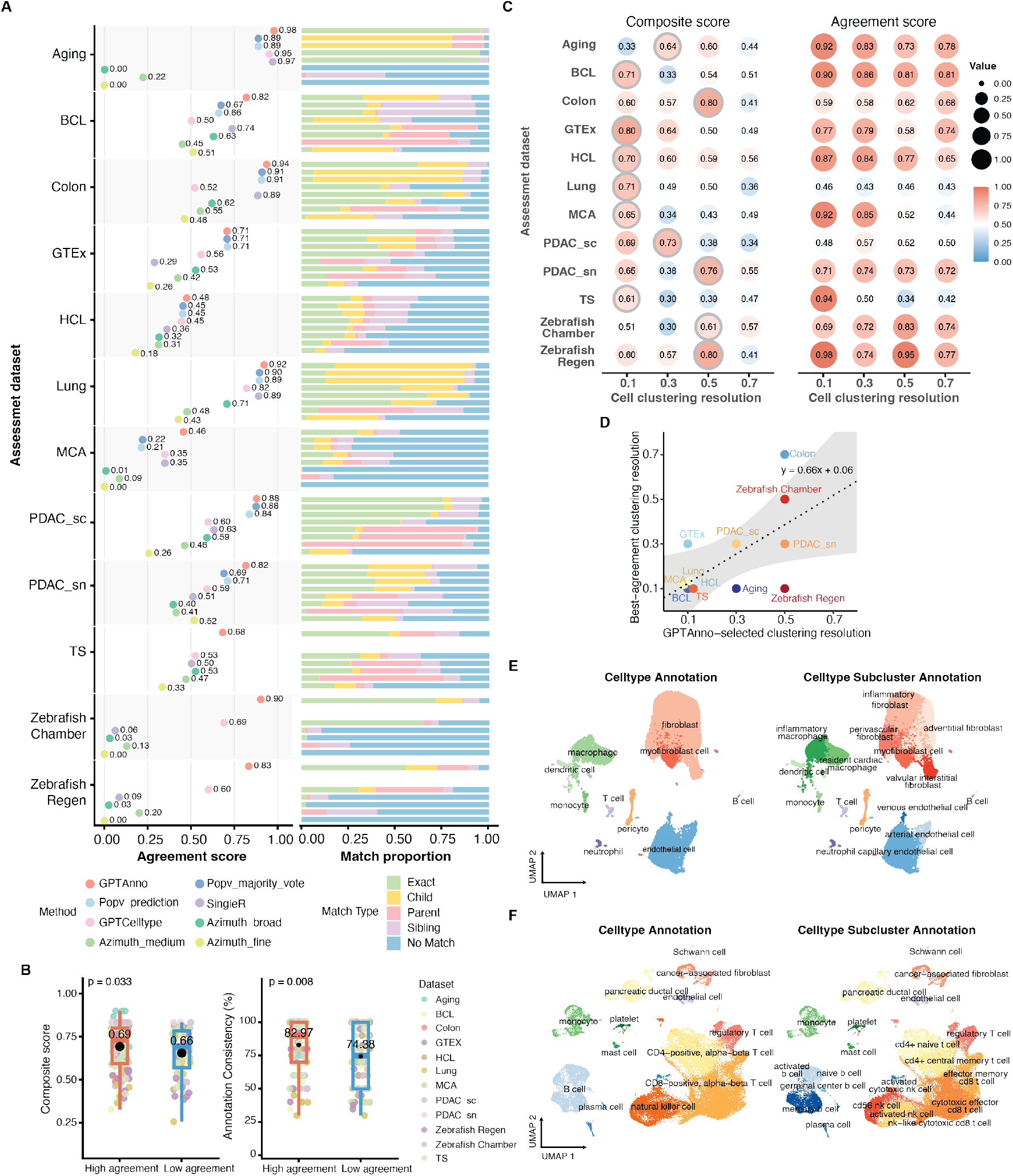
Systematic evaluation of GPTAnno across 12 datasets. **A)** Agreement scores and decomposed matches between predicted and expert labels. Note: Tabula Sapiens (TS) does not include popV predictions, as popV was pretrained on TS. **B)** Boxplots showing high-agreement (agreement > 0.5) and low-agreement (agreement ≤ 0.5) datasets against composite scores (left) and annotation consistency (right). **C)** Multi-resolution performance across benchmark datasets. Circled values indicate the largest values that led GPTAnno to select resolutions. **D)** Scatterplot of clustering resolution selected by GPTAnno versus by best-agreement. Dotted line indicates linear fitting with grey area indicating 95% confidence interval. **E & F)** Case study of mouse heart aging (E) and PDAC_sc (F) data hierarchical annotation. Left: broad cell type annotation at the optimal resolution selected by GPTAnno. Right: Cell type subcluster annotation by marker inheritance strategy.

To examine how GPTAnno’s performance relates to the composite scores it used for choosing clustering resolution, we split assessment datasets into high-agreement (> 0.5, closely matching expert labels) and low-agreement (≤ 0.5) groups. The high-agreement group showed higher composite scores (mean 0.69) and higher annotation consistency (mean 82.97%) than the low-agreement group (mean 0.66 and 74.38% separately), respectively (Fig. 2B). Two-sided Wilcoxon rank sum tests confirmed these differences (composite score *p* = 0.033, consistency *p* = 0.008). To visualize how composite and agreement scores vary across clustering resolutions, we reclustered each dataset at 0.1, 0.3, 0.5, and 0.7 and then evaluated agreement with expert labels. At each clustering resolution, we ran the remaining GPTAnno steps and evaluated the agreement with the original expert labels. Note that GPTAnno’s automated pipeline selected the resolution with the largest composite score. In eight out of twelve datasets, the selected resolution delivered the highest agreement. In the remaining four (Aging, GTEx, PDAC single-nucleus, Zebrafish Regeneration), agreement at the selected resolution was within 10% of the best alternative (Fig. 2C, Supplementary Table 2). To avoid favoring any fixed resolution by including datasets where that resolution is also selected by GPTAnno, we show a variant where datasets are excluded from a fixed-resolution box if GPTAnno also selected that resolution (Supplementary Fig. S2). Overall, adaptive resolution via GPTAnno typically matched the best-agreement setting, reducing trial-and-error repeated re-clustering and underscoring the value of the ontology-aware selection step (Fig. 2D).

To illustrate the use of hierarchical clustering and its flexibility, we performed case studies on two datasets that were published beyond GPT-5’s (API version) training cutoff date: mouse heart aging data^46^ and PDAC single-cell RNA-seq data^48^(Fig. 2E, F). Following broad-level annotation, we applied GPTAnno’s marker inheritance strategy to refine cell populations: for the PDAC dataset, we subclustered CD4-positive alpha-beta T cells, CD8-positive alpha-beta T cells, B cells, and natural killer cells; for the aging dataset, we subclustered endothelial cells, fibroblasts, and macrophages. The annotations were validated through comparison with marker genes and consultation with domain experts (Supplementary Figure S1).

To further explore the potential for ontology enhancement, we developed an AI literature mining module that automatically extracts and summarizes emerging cell types from scientific articles. Ten relevant papers were selected, and their PDF files were processed through the pipeline. The outputs (Supplementary Table 3) include both the final extraction results of cell types and marker genes mentioned in the paper and example intermediate parsing outputs from ten single-cell cardiac neonatal research. This module is a standalone literature-mining tool extensively enhancing GPTAnno’s ontology-based annotation strategy, allowing automated integration of newly described cell populations into GPTAnno’s framework.

Beyond benchmarking accuracy, our composite scoring framework proved highly informative, not only quantifying performance during prediction but also revealing annotation inconsistencies and biological ambiguities. The prediction with high consistency during prediction but low agreement during evaluation could indicate potential errors in manual labels. For example, in the BCL dataset, GPTAnno confidently annotated one cluster as plasma cells, supported by strong markers such as *JCHAIN*(joining chain of IgA/IgM) and *IGHG4* (immunoglobulin heavy constant gamma 4). Although the original study labeled this cluster as B cells, our scoring framework flagged the discrepancy, suggesting that the plasma cell annotation may be more biologically accurate. In addition, GPTAnno can highlight the mixed cells or ambiguous cell types. In aging and MCA datasets, GPTAnno repeatedly predicts one cluster as smooth muscle cell (CL:0000192) or pericyte (CL:0000669, approximately 50%/50% across runs), which is a biologically plausible ambiguity given their spatial proximity, shared markers, and the possibility of RNA contamination^50^. The manual labels also annotated these clusters as mixed, further validating the framework’s diagnostic power.

Taken together, our ontology-aware scoring is not only evaluative but also a discovery tool. In doing so, GPTAnno provides a rigorous, ontology-based lens for assessing annotation accuracy, supporting consistency, and revealing lineage coherence across diverse datasets.

## 3 Methods

### 3.1 Preprocess and Multi-resolution Clustering

Raw single-cell RNA-seq expression matrices were optionally processed using the Seurat (v5.2.0) package^51^. The `preprocess_seurat_object` function normalized gene expression values (LogNormalize, scale factor: 10,000), identified highly variable features (default: 3,000), scaled the data, and performed principal component analysis (PCA) using the top variable genes. Cell clustering was performed using the `run_multi_resolution_clustering` function, which performed graph-based clustering across multiple candidate resolutions defined by the user (default range: 0.1 to 0.7 in increments of 0.2). At each resolution, clusters were identified using Louvain algorithm^52^. Differential expression analysis was performed using the Wilcoxon rank-sum test to identify marker genes for each cluster, filter genes with log_2_-fold-change > 1, and select the top *n* genes (default: *n* = 10)^26^. Marker gene lists and cluster assignments were saved for each resolution to construct the prompt.

### 3.2 LLM Query and Cell Type Annotation

For each clustering resolution, the marker genes with the smallest *p*-values (default: top 10) and, if ties, largest absolute difference in the percentage of cells expressing the gene between clusters were selected to construct the inputs to the `gptcelltype` function in GPTCelltype package^26^. The GPTCelltype^26^ prompting strategy has the following template: *“Identify cell types of [TissueName] cells using the following markers separately for each row. Only provide the cell type name. Do not show numbers before the name. Some can be a mixture of multiple cell types. [GeneList]”*. Here, [TissueName] is replaced with the provided tissue context (e.g., human prostate), and [GeneList] contains marker genes for each cluster separated by newlines, with genes within clusters separated by commas. All LLM queries in the evaluation used OpenAI’s GPT-5 model (gpt-5-2025-08-07) accessed via the OpenAI API. API Query was through openai (v0.4.1) R package. To quantify LLM prediction variability, GPTAnno performs repeated queries per resolution (default: 10). Both the model and the number of queries can be specified by the user. Raw LLM outputs of cell type predictions are systematically cleaned and mapped to CL using the `clean_and_match_annotation` function. This multi-step process includes: (1) string normalization (lowercasing, whitespace trimming, singularization, removal or addition of common suffixes like “cells”), (2) synonym substitution using a curated mapping table `GPTCelltype_mapping`, which maps GPT style annotations to standardized CL names (curated from Hou et al.^26^, e.g., “cardiomyocyte” to “cardiac muscle cell”), and (3) mapping using `cl_term_map`, which matches names to official CL identifiers with keys or synonyms (generated from `build_cl_term_map`). If no direct match is found, users can optionally enable an Ontology Lookup Service (OLS) search^53^ with `search_ols`. This search returns top-ranked CL terms based on word overlap and semantic similarity, providing suggested ontology mappings for ambiguous or novel predictions.

### 3.3 Optimal Number of Clusters Selection

We developed a composite scoring framework to identify the clustering resolution that yields the most biologically coherent annotations. Consider candidate resolutions {*r*_1_, …, *r*_*K*_}, where resolution *r*_*k*_ ∈ (*k* [1, *K*]) produces *C*_*k*_ clusters 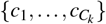. Each query round *t* = 1, …, *R* returns predictions for all clusters at that resolution. Denote the raw prediction for cluster *c* in query *t* as *y*_*c,t*_, its standardized prediction (CL term) as 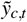, and the total number of unique CL terms in cluster *c* as *m*_*c*_. For each cluster *c*, the selected annotation is the prediction with the highest frequency across the *R* queries. The corresponding maximum frequency (percentage) is denoted 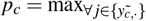 Pr(annotation = *j* | cluster *c*). Using these values, three resolution-level metrics are defined:

1. consistency, the average of maximum prediction frequency across clusters, 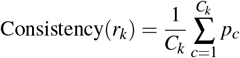
2. robustness, the minimum of the maximum frequencies, which identifies the weakest (least stable) cluster in resolution *r*_*k*_;

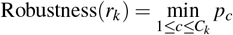
3. reliability, the average ontology dispersion across clusters. For each cluster *c*, dispersion is computed as the mean pairwise shortest-path distance on the ontology tree among its unique standardized predictions,

#### Algorithm 1

Composite Scoring for Resolution Selection in GPTAnno

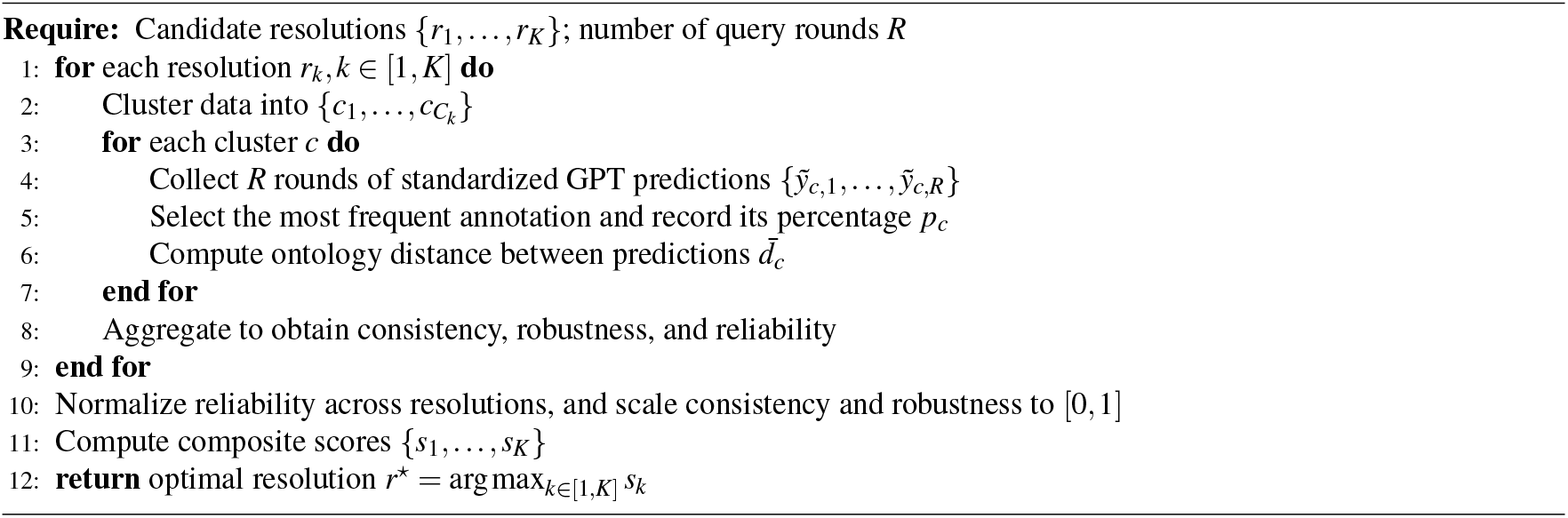

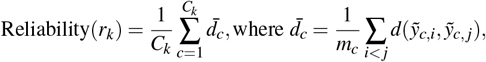 The reliability metric is normalized across resolutions and inverted (1 − normalized distance), so that higher values correspond to greater reliability; percentages (consistency and robustness) are scaled directly to [0, 1]. The composite score for resolution *r* _*j*_ is then

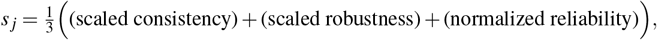

and the optimal resolution is selected as *r*^⋆^ = argmax_*k*_ *s*_*k*_.

### 3.4 Multi-resolution Subclustering and Hierarchical Annotation

Hierarchical annotation is routinely used by human experts to refine broad categories into more specific subtypes. The Cell Ontology (CL) encodes such hierarchical relationships and can guide subcluster annotation, though many emerging or transitional cell types described in recent literature are not yet represented. GPTAnno first identifies which parent cell types should be further annotated as the subtypes (children cell types). Users may directly specify the cell types of interest for subclustering. Alternatively, GPTAnno can automatically identify candidates based on criteria: cell type that has descendants on CL and exceeds a minimum cell count (default: 10,000). For each selected parent type, we developed two prompting strategies tailored for hierarchical annotation. *1. CL-restricted prompting*. When a parent cell type is represented in CL, the LLM prompt is explicitly restricted to the child terms of that parent on the ontology tree. For example, if the parent cluster is annotated as “T cell,” the model is instructed to select among its CL-defined descendants (e.g., CD4-positive T cell, CD8-positive T cell). This approach enforces consistency with ontology-defined hierarchies and prevents the model from assigning unrelated labels. *2. Parent marker inherited prompting*. As some cell types may not have descendants or those descendants are not relevant to the study, the prompt would add the parent cluster’s marker genes to the subcluster’s top marker genes, without restricting to any CL terms. The parent markers provide general lineage context (e.g., canonical T cell markers), while the subcluster markers highlight distinguishing features (e.g., cytotoxicity-related genes). By combining both, the model can propose biologically coherent subtypes, leveraging the flexibility of LLMs, even when they are absent from CL. As with parent-level annotation, GPTAnno evaluates multiple subcluster resolutions and selects the optimal result using the same composite scoring framework that combines consistency across repeated queries with reliability derived from ontology distances.

### 3.5 AI-Assisted Literature Mining for Novel Cell Types

To systematically identify novel cell types and their associated marker genes from published literature, we developed an AI-assisted literature mining module, PDF2Markers. The pipeline integrates PDF text extraction, LLM-based entity recognition powered by `gpt-5-nano`, and ontology-guided filtering to produce high-consistency, structured annotations suitable for seamless integration into the GPTAnno framework.

Scientific articles in PDF format were parsed via `pypdf`^54^ to extract text across all pages. To obtain publication metadata, the pipeline analyzes the text from the first page and applies a GPT-based parser (`gpt-5-nano`) that is prompted to identify and return structured fields—including the first author’s surname, journal name, and publication year—in JSON format. When GPT parsing is inconclusive, a fallback heuristic extracts the publication year via regular-expression matching. The full text was segmented into sentences and grouped into short overlapping windows to preserve local context. Candidate sentences containing marker-related keywords (e.g., marker, expressed, enriched) were identified and analyzed by GPT models through structured prompts that requested JSON-formatted cell type–marker gene pairs. To ensure robustness and efficiency, the model queries were executed in parallel batches with on-disk caching of prior responses.

Extracted results were subsequently subjected to a series of post-processing and quality-control procedures. Abbreviated or short-form cell-type names detected in the LLM outputs were first expanded to full names using two complementary sources: (i) abbreviation–full-name mappings automatically mined from the PDF (e.g., patterns such as “Full Name (Short)”, and (ii) domain-specific heuristic rules encoding common biological abbreviations (see Table 1). After expansion, all cell-type names underwent standardized normalization, including lowercasing, whitespace collapsing, removal of trivial morphological variants (e.g., cells → cell), and rule-based singularization to harmonize plural forms. Additional cleanup steps filtered out ambiguous or non-biological labels, such as names containing “-like” or patterns beginning with a letter–digit prefix. The resulting normalized and validated full-length names were then used for downstream deduplication and marker-gene consolidation. All cleaned outputs were saved in standardized CSV files with provenance information and quality-control flags. The full source code, including modules for document retrieval, PDF parsing, language-model extraction, and post-processing integration, is publicly available in the `GPTAnno/PDF2markers` repository.

### 3.6 Dataset collection

We used 12 datasets from 10 studies, including Genotype-Tissue Expression (GTEx)^39^, Human Cell Landscape (HCL),^40^, Mouse Cell Atlas (MCA),^41^, Tabula Sapiens (TS)^42^, B-cell lymphoma^43^, lung cancer^45^, colon cancer^44^, mouse aging heart^46^, zebrafish heart injury and regeneration^47^, and PDAC tumor and adjacent normal tissues^48^ datasets. The last two papers provide two datasets separately. For datasets containing multiple modalities or conditions, we treated each as a separate dataset. The zebrafish study provides two scRNA-seq datasets: (1) uninjured adult chamber and (2) pre- and post-injury heart regeneration. The PDAC study provides: (1) scRNA-seq from tumor and adjacent normal tissues, and (2) snRNA-seq from normal pancreatic tissue only. For each dataset, we downloaded the gene-by-cell expression matrix (count matrix, RDS file, or h5ad file) and the author-provided manual cell type annotations directly from the publication supplement or linked repositories. All scRNA-seq datasets were processed using a standardized pipeline. Following clustering, differentially expressed genes were identified for every cluster using Seurat’s `FindAllMarkers`. For HCL and MCA, which profile dozens of tissues, cell type annotation and evaluation were performed once per atlas by aggregating all tissues into a single object, mirroring the approach in the original studies. For the PDAC dataset, which combines normal, tumor, and peripheral blood mononuclear cells (PBMCs) tissues, we first integrated normal and cancer cells using canonical correlation analysis (CCA) via Seurat’s `IntegrateLayers`. This integration produces a shared low-dimensional representation that aligns biological structure across conditions and serves as the basis for downstream unsupervised clustering and marker detection.

### 3.7 External Annotation Methods

To benchmark GPTAnno against representative external methods, we applied GPTCelltype^26^, popV^15^, AzimuthAPI^16^ and SingleR^11^, using standardized preprocessing across all datasets.

#### SingleR

We used the `SingleR` package (v2.8.0) with the following wrapper:

~~~
annotate_with_SingleR <- function(seurat_object, ref) {
   # Convert to SingleCellExperiment
   sce <- as.SingleCellExperiment(seurat_object)
   # Run SingleR annotation
   predictions <- SingleR(test = sce, ref = ref, labels = ref$label.main)
   seurat_object$SingleR_labels <- predictions$labels
   return(list(seurat_object = seurat_object, metadata = seurat_object@meta.data))
}
~~~

Reference atlases (e.g., Human Primary Cell Atlas and Tabula Sapiens) were used as in the original SingleR publication.

#### AzimuthAPI

We used the AzimuthAPI (v0.1.0) R package to interface directly with Seurat’s cloud-based Azimuth service:

~~~
annotate_with_Azimuth <- function(seurat_object){
   library(AzimuthAPI)
   # Run annotation (cloud-based R API)
   seurat_object <- CloudAzimuth(seurat_object)
   # Extract predictions
   meta <- seurat_object@meta.data
   seurat_object$Azimuth_final <- meta$final_level_labels
  seurat_object$Azimuth_medium <- meta$azimuth_medium
   seurat_object$Azimuth_broad <- meta$azimuth_broad
   return(list(seurat_object = seurat_object))
}
~~~

Azimuth outputs (*broad, medium*, and *fine*) were imported into the Seurat metadata.

#### popV

The `popV` method was implemented following the official workflow from the Yosef Lab repository (https://github.com/YosefLab/popV). We used the default ensemble configuration to generate both popV majority vote and popV prediction annotations.

#### GPTCelltype

We also compared to `GPTCelltype` (https://github.com/Winnie09/GPTCelltype), which uses GPT based marker interpretation. For each dataset, marker genes derived from the GPTAnno-selected optimal resolution were supplied to the `gptcelltype()` function using the `gpt-5 (gpt-5-2025-08-07)` model. For consistency, we executed three independent runs per dataset and retained the prediction achieving the highest agreement score for downstream evaluation, to minimize stochastic variation in LLM outputs. Note that our ontology-aware evaluation uses weighted agreement scores based on CL hierarchy (detailed in the next section), while the original study used exact match, partial match, and no match to calculate annotation agreement. This difference may result in higher or lower apparent performance, depending on whether predictions are more specific or more general than manual labels.

### 3.8 Evaluations of cell type annotations

All annotations were evaluated using ontology-aware function (`score_annotation_agreement_ontology_detailed`) to quantify hierarchical correspondence (*Exact, Parent, Child, Sibling, No Match*) relative to manual curation. For each annotated Seurat object, both manual and predicted labels were mapped to CL term IDs using the built-in `cl_term_map` reference. Ontology relationships were derived from the Cell Ontology API (http://purl.obolibrary.org/obo/cl.obo), loaded with `ontologyIndex::get_ontology()` and represented as a directed acyclic graph via `igraph`. Each pair of manual and predicted annotations was classified into one of five hierarchical relationship types: **Exact** (same CL ID), **Parent** (predicted is an ancestor of the manual label), **Child** (manual is an ancestor of the predicted label), **Sibling** (distance ≤ 3 on the CL tree with shared common ancestors but no direct ancestry), or **No match** (no detectable relationship within the ontology). Each match type was assigned a weighted score (exact = 1.0, child = 1.0, parent = 0.5, sibling = 0.5, no_match = 0.0) to generate an overall agreement score per cell. Summary statistics were then computed across all cells, including mean agreement, frequency of each match type, and ontology distance distributions. For transparency and reproducibility, per-cell detailed results, summary statistics, and pairwise match tables were exported.

### 3.9 API Query Cost

Each GPTAnno annotation task was executed for three clustering resolutions, with ten repeated GPT queries per resolution to ensure reproducibility (30 queries per dataset). Using OpenAI’s GPT-5 pricing structure ($1.25/1M input tokens, $0.125/1M cached input tokens, and $10.00/1M output tokens), each query with a total of 1,000–5,000 tokens cost approximately $0.0017–$0.0084. This corresponds to an estimated cost of $0.05–$0.25 per dataset. Similarly, for subclustering, the total token cost depends on the number of cell types selected for subclustering and the number of candidate resolutions evaluated per cell type. Datasets with more subclustered cell types or additional resolution will have higher query counts, though the per-query cost remains within the same range.

## 4 Discussion

In this study, we developed an automated framework for scRNA-seq cell type annotation that integrates LLM with CL to find the optimal clustering resolution and provide accurate predictions. Beyond being powered by the LLM for tools like GPTCelltype, GPTAnno enables systematic evaluation of LLM-based predictions for consistency and reliability, supports hierarchical annotation, and extends the current CL by LLM’s own ability and mining the literature for up-to-date cell type definitions. Our study has several limitations. First, we observed that GPTAnno’s predictions tend to favor smaller resolutions with fewer clusters. This highlights the need for hierarchical annotation, where broad categories can be further subdivided into biologically coherent subtypes. Second, we note that marker-based predictions could be influenced by batch effects, which can bias differential gene expression and downstream LLM prompts. We advise users to perform batch correction to improve robustness. Third, the pipeline inherits the double-dipping issue common in clustering workflows, as clustering and differential gene expression are both performed on the same dataset. Selective inference approaches (e.g., Kmeansinference^55^) could be options to mitigate these problems. Finally, we emphasize our effort to use CL as the backbone for standardization and evaluation. CL greatly improves consistency across queries and enables ontology-aware scoring, yet its own limitations remain: some cell types are missing, others are defined inconsistently, and many recent or marker-defined populations have not yet been incorporated. Attention to these ontology gaps is critical, and ongoing updates to CL will be essential for the continued success of ontology-guided annotation.

## Declarations

### Code availability

The full implementation of `GPTAnno`, including clustering, ontology-based reasoning, and scoring modules, is openly available at: https://github.com/yrsong001/GPTAnno. All scripts were implemented in `R` (version ≥ 4.2) and `Python` (version ≥ 3.8) using standard open-source dependencies.

## Acknowledgments

W.H. was supported by the National Institutes of Health (NIH) under Award Number R00HG011468 (NHGRI) and R35GM150887 (NIGMS). F.Z. was supported by the NIH under Award Number R01HL173044, R01LM014407, and P42 ES031007. Y.S. is supported by NIH R35HL155656 (NHLBI), P42 ES031007. Q.L. was supported by the NIH under Award number R00HG011468 (NHGRI). L.Q. is supported by NIH R35HL155656 (NHLBI) and AHA 20EIA35320128. H.W. is supported by AHA 23CDA1042496.

## Author contributions

W.H. and F.Z. conceived the study. Y.S., W.H., and F.Z. developed the method. Y.S., M.T., Q.L., and W.H. performed the analysis. H.W. and L.Q. provided biological interpretation and domain expertise. All authors contributed to the writing of the manuscript.

## A Supplement Figure

## B Supplement Table

**Table S1.** Comparison of GPTAnno with existing single-cell annotation methods. GPTAnno uniquely combines reference-free clustering, Cell Ontology standardization, reproducibility quantification, and LLM-assisted reasoning. ✓ = supported; ✗ = not supported; △ = conditional/depends on tools invoked.

| Feature | GPTAnno | GPTCelltype | SingleR | Azimuth | PopV | CellTypist | Transformer<br>(scGPT,<br>scBERT) | scExtract | Agent-based<br>(CellAgent,<br>scAgent) |
| --- | --- | --- | --- | --- | --- | --- | --- | --- | --- |
| *Annotation Ability* |  |  |  |  |  |  |  |  |  |
| Reference-free annotation | ✓ | ✓ | ✗ | ✗ | ✓ | ✗ | ✓ | ✓ | △ |
| Ontology standardization | ✓ | ✓ | ✗ | ✗ | ✓ | ✗ | ✗ | ✗ | ✗ |
| Optimal resolution selection | ✓ | ✗ | ✗ | ✗ | ✗ | ✗ | ✗ | ✗ | ✗ |
| Hierarchical subclustering | ✓ | ✗ | ✗ | ✗ | ✓ | ✗ | ✗ | ✗ | ✗ |
| Multi-level label provision | ✓ | ✗ | ✗ | ✓ | ✓ | ✗ | ✗ | ✗ | △ |
| *Quality assurance* |  |  |  |  |  |  |  |  |  |
| Reproducibility quantification | ✓ | ✗ | ✗ | ✗ | ✗ | ✗ | ✗ | ✗ | ✗ |
| Ontology distance metrics | ✓ | ✗ | ✗ | ✗ | ✗ | ✗ | ✗ | ✗ | ✗ |
| Prediction consistency scoring | ✓ | ✗ | ✗ | ✗ | ✗ | ✗ | ✗ | ✓ | ✓ |
| *Ontology integration* |  |  |  |  |  |  |  |  |  |
| Standardized CL term mapping | ✓ | ✓ | ✗ | ✗ | ✓ | ✗ | ✗ | ✗ | ✗ |
| CL graph based evaluation | ✓ | ✗ | ✗ | ✗ | ✓ | ✗ | ✗ | ✗ | ✗ |
| *Advanced capabilities* |  |  |  |  |  |  |  |  |  |
| General-purpose LLM integration | ✓ | ✓ | ✗ | ✗ | ✗ | ✗ | ✗ | ✓ | ✓ |
| Novel type literature mining | ✓ | ✗ | ✗ | ✗ | ✗ | ✗ | ✗ | ✓ | △ |
| Seurat compatibility | ✓ | ✓ | ✓ | ✓ | ✗ | ✗ | ✓ | ✗ | △ |

**Figure S1.**
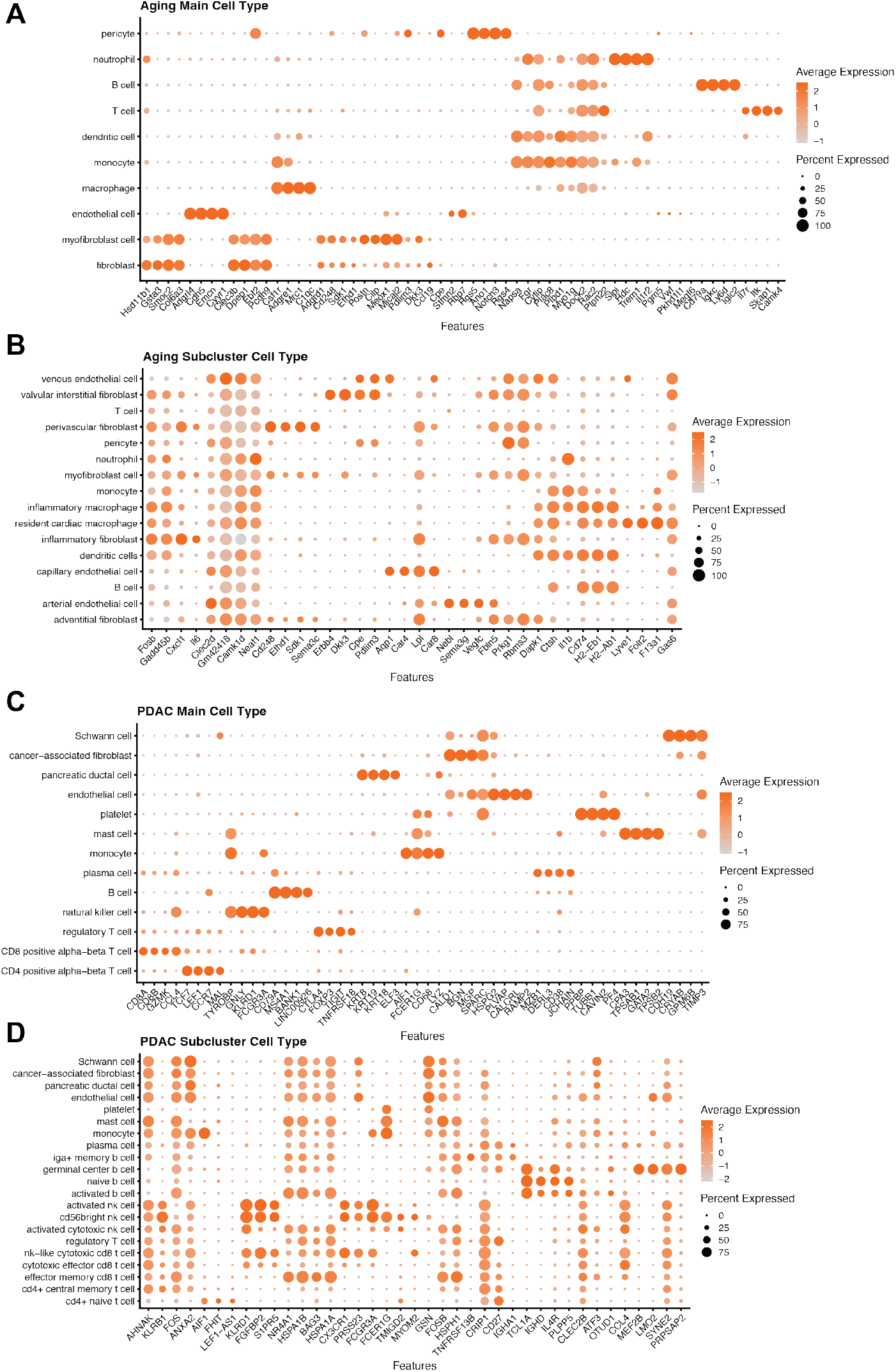
Marker gene validation for hierarchical cell type annotation. **A)** Dot plot showing marker gene expression patterns for main cell types identified in the mouse heart aging dataset. Dot size represents the percentage of cells expressed. Color intensity indicates average expression level. **B)** Marker gene expression for refined subcluster cell types in the aging dataset following GPTAnno’s hierarchical annotation. **C)** Dot plot showing marker gene expression patterns for main cell types identified in the PDAC single-cell dataset. **D)** Marker gene expression for refined subcluster cell types in the PDAC dataset.

**Figure S2.**
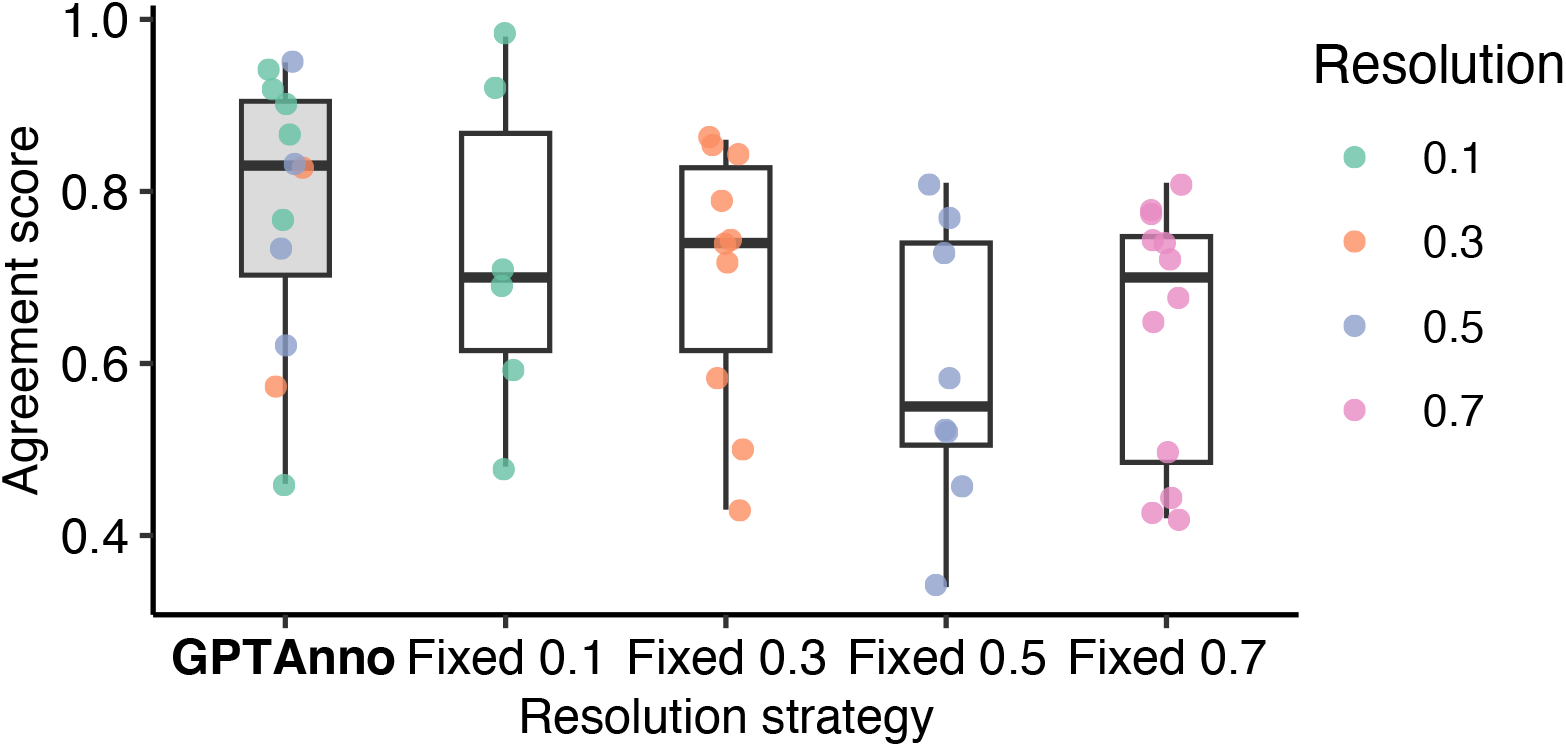
Comparison of GPTAnno’s adaptive resolution selection strategy versus fixed-resolution approaches. Boxplots show agreement score distributions across datasets for optimal resolution selection (GPTAnno) and fixed resolutions (0.1, 0.3, 0.5, 0.7).

**Table S2.** Ablation analysis of the relationship between the chosen resolution (selected by composite scoring) and the ontology-based agreement score across twelve datasets.

| Dataset | Resolution | Agreement Score | Composite Score | Avg Path Length |
| --- | --- | --- | --- | --- |
| Aging | 0.1 | 0.979 | 0.601 | 1.407 |
|  | 0.3 | 0.737 | 0.569 | 1.231 |
|  | **0.5** | **0.950** | **0.797** | **0.979** |
|  | 0.7 | 0.771 | 0.407 | 1.553 |
| BCL | 0.1 | 0.689 | 0.508 | 1.667 |
|  | 0.3 | 0.723 | 0.298 | 1.825 |
|  | **0.5** | **0.835** | **0.610** | **1.387** |
|  | 0.7 | 0.743 | 0.574 | 1.452 |
| Colon | **0.1** | **0.937** | **0.606** | **0.385** |
|  | 0.3 | 0.500 | 0.298 | 0.717 |
|  | 0.5 | 0.344 | 0.385 | 0.600 |
|  | 0.7 | 0.422 | 0.473 | 0.506 |
| GTEx | 0.1 | 0.708 | 0.652 | 0.617 |
|  | 0.3 | 0.744 | 0.383 | 0.744 |
|  | **0.5** | **0.726** | **0.760** | **0.563** |
|  | 0.7 | 0.719 | 0.553 | 0.667 |
| HCL | 0.1 | 0.475 | 0.692 | 0.708 |
|  | **0.3** | **0.567** | **0.729** | **0.640** |
|  | 0.5 | 0.524 | 0.382 | 0.953 |
|  | 0.7 | 0.496 | 0.344 | 0.930 |
| Lung | **0.1** | **0.922** | **0.646** | **0.841** |
|  | 0.3 | 0.851 | 0.339 | 1.270 |
|  | 0.5 | 0.517 | 0.431 | 1.116 |
|  | 0.7 | 0.443 | 0.489 | 0.955 |
| MCA | **0.1** | **0.456** | **0.706** | **0.498** |
|  | 0.3 | 0.432 | 0.486 | 0.648 |
|  | 0.5 | 0.456 | 0.501 | 0.653 |
|  | 0.7 | 0.435 | 0.358 | 0.801 |
| PDAC_sc | **0.1** | **0.869** | **0.700** | **0.462** |
|  | 0.3 | 0.842 | 0.602 | 0.500 |
|  | 0.5 | 0.767 | 0.585 | 0.514 |
|  | 0.7 | 0.647 | 0.557 | 0.508 |
| PDAC_sn | **0.1** | **0.768** | **0.798** | **0.300** |
|  | 0.3 | 0.789 | 0.641 | 0.476 |
|  | 0.5 | 0.579 | 0.498 | 0.889 |
|  | 0.7 | 0.741 | 0.491 | 0.789 |
| Zebrafish Chamber | **0.1** | **0.902** | **0.706** | **1.071** |
|  | 0.3 | 0.857 | 0.334 | 1.477 |
|  | 0.5 | 0.805 | 0.536 | 1.237 |
|  | 0.7 | 0.815 | 0.506 | 1.244 |
| Zebrafish Regeneration | 0.1 | 0.919 | 0.327 | 1.786 |
|  | **0.3** | **0.834** | **0.636** | **1.345** |
|  | 0.5 | 0.731 | 0.602 | 1.487 |
|  | 0.7 | 0.777 | 0.438 | 1.552 |
| TS | 0.1 | 0.590 | 0.601 | 1.407 |
|  | 0.3 | 0.581 | 0.569 | 1.231 |
|  | **0.5** | **0.624** | **0.797** | **0.979** |
|  | 0.7 | 0.683 | 0.407 | 1.553 |

